# HIV-1 subtype C with PYxE insertion has enhanced binding of Gag-p6 to host cell protein ALIX and increased replication fitness

**DOI:** 10.1101/459156

**Authors:** Robert van Domselaar, Duncan T. Njenda, Rohit Rao, Anders Sönnerborg, Kamalendra Singh, Ujjwal Neogi

**Author notes:** These authors contributed equally (Robert van Domselaar, Duncan T. Njenda). **Correspondence:** Ujjwal Neogi, Division of Clinical Microbiology, Department of Laboratory Medicine, Karolinska Institutet, Stockholm, Sweden, e-mail.

## Abstract

Human immunodeficiency virus type 1 subtype C (HIV-1C) has a natural deletion of a YPxL motif in its Gag-p6 late domain. This domain mediates the binding of Gag to host cell protein ALIX and subsequently facilitates viral budding. In a subset of HIV-1C infected individuals, the tetrapeptide insertion PYxE has been identified at the deleted YPxL motif site. Here, we report the consequences of PYxE insertion on the interaction with ALIX and the relevance regarding replication fitness and drug sensitivity. In our three HIV-1C cohorts, PYKE and PYQE were most prevalent among PYxE variants. Through *in silico* predictions and *in vitro* experiments, we showed that HIV-1C Gag has an increased binding to ALIX when PYxE motif is present. To go more into the clinical relevance of the PYxE insertion, we obtained patient-derived gag-pol sequences from HIV-1C_PYxEi_ viruses and inserted them in a reference HIV-1. Viral growth was increased, and the sensitivity to protease inhibitor (PI) lopinavir (LPV) and nucleoside reverse transcriptase inhibitor tenofovir alafenamide (TAF) was decreased for some of the HIV-1C PYxE variants compared to wild-type variants. Our data suggest that PYxE insertion in Gag restores the ability of Gag to bind ALIX and correlates with enhanced viral fitness in the absence or presence of LPV and TAF. The high prevalence and increased replication fitness of the HIV-1C virus with PYxE insertion could indicate the clinical importance of these viral variants.

**Importance:** Genomic differences within HIV-1 subtypes is associated with a varying degree of viral spread, disease progression, and clinical outcome. Viral budding is essential in the HIV-1 life cycle and mainly mediated through the interaction of Gag with host proteins. Two motifs within Gag-p6 mediate binding of host cell proteins and facilitate budding. HIV-1 subtype C (HIV-1C) has a natural deletion of one of these two motifs resulting in an inability to bind to host cell protein ALIX. Previously, we have identified a tetrapeptide (PYxE) insertion at this deleted motif site in a subset of HIV-1C patients. Here, we report the incidence of PYxE insertions in three different HIV-1C cohorts, and the insertion restores the binding of Gag to ALIX. It also increases viral growth even in the presence of antiretroviral drugs lopinavir and tenofovir alafenamide. Hence, PYxE insertion in HIV-1C might be biologically relevant for viruses and clinically significant among patients.

## Introduction

Human immunodeficiency virus type 1 (HIV-1) is a global threat with an estimated 36.9 million HIV-1 infected individuals and 940,000 deaths in 2017. Virulence of HIV-1 is determined by its capacity to replicate within infected cells and its ability to infect new cells. Among different HIV-1 subtypes, more than 50% of all infections are caused by HIV-1 subtype C (HIV-1C) that is prevalent in South Africa, Zimbabwe, Mozambique, Botswana, Ethiopia, Eritrea, India, and in some parts of Brazil. However, HIV-1C has also become highly prevalent in several European countries, including Sweden (1, 2). The *gag* gene encodes for a polyprotein that is proteolytically processed by viral and cellular proteases into six final products: p17 matrix protein (MA), p24 capsid protein (CA), spacer peptide 1 (SP1), p7 nucleocapsid protein (NC), spacer peptide 2 (SP2), p6 protein (P6). These proteins are required for the assembly and release of new virions (3-6). The p6 late domain of Gag contains two conserved short peptide motifs that bind host factors to facilitate viral budding (6-8). The PTAP motif is present in both HIV-1 and HIV-2, and sometimes it is duplicated in HIV-1 genomes (9, 10). It mediates binding to ESCRT-I subunit tumor susceptibility gene 101 (TSG101) (11). The YPx_n_L (where x_n_ represents a random number of any possible amino acids) motif binds to ESCRT-III adaptor protein ALG-2-interacting protein X (ALIX) (12). However, this latter motif is deleted exclusively in HIV-1C (13). This deletion is associated with the loss of binding of p6 late domain to ALIX and a decrease in virus release from infected cells. Of note, HIV-2 lacks the YPx_n_L motif but contains an alternative ALIX-binding motif, namely PYKEVTEDL, that originates from simian immunodeficiency virus (SIV) in rhesus macaques (14).

Recently, we identified a tetrapeptide insertion PYxE [where x represents lysine (K), glutamine (Q) or arginine (R)] within the Gag protein in a subgroup of HIV-1C infected individuals (C_PYxEi_-strains) (15). This C_PYxEi_-strain was preferentially identified in East African HIV-1C infected patients but was less common among HIV-1C infected patients from South Africa, India, and Germany (15). Furthermore, the insertion appears more frequent among patients on antiretroviral therapy, e.g., ritonavir-boosted protease inhibitors (PI), compared to therapy naïve patients in India, and more frequent in therapy-failure patients in South Africa (15, 16). Moreover, lower pre-therapy CD4^+^ T cell counts, higher plasma viral loads, and reduced increase in CD4^+^ T cell counts were noted to be associated with PYxE insertion in HIV-1C patients compared to patients with wild type HIV-1C from East Africa (17). Furthermore, we observed that increased replication fitness in PYQE inserted HIV-1C viruses was polymerase independent (17). A more recent study also claimed that gag-protease is the major determinant of subtype differences in disease progression among HIV-1 subtypes (18).

As the tetrapeptide PYxE insertion was found at the site of the lost YPx_n_L motif of HIV-1C Gag-p6, we hypothesized that this PYxE insertion might restore the interaction of the Gag-p6 late domain with ALIX and thereby increases replication fitness and reduces sensitivity to PIs. A recent report showed that the PYRE insertion could rescue the viral growth of a PTAP-deleted HIV-1 variant similar to the YPx_n_L motif in the absence of a PTAP sequence (19). However, since the PTAP motif is always present in clinical HIV-1 isolates, it is not known if and how PYxE insertion affects HIV-1C pathogenicity in a clinically relevant context. To address this, we assessed the distribution and frequency of this PYxE insertion in different cohorts of HIV-1C infected individuals. We also evaluated the ability of PYxE motif to reconstitute the interaction of Gag with ALIX, its association with increased viral growth of clinical isolates, and its association with drug responses against all three drug classes; reverse transcriptase inhibitors (RTIs), PIs, and integrase strand transfer inhibitors (INSTIs).

## Material and Methods

### Cell culture and plasmids

Cells were cultured in 5% CO_2_ at 37°C. HEK293T cells were maintained in Dulbecco’s modified Eagle medium (DMEM, Sigma, USA) supplemented with 10% fetal calf serum (Sigma,USA), 2 mM L-glutamine (Sigma, USA), 0.1 mM MEM Non-Essential Amino Acids (Gibco/Thermo Fisher Scientific, USA), and 10 units/mL penicillin combined with 10 μg/mL streptomycin (Sigma). TZM-bl reporter cells were also maintained in medium mentioned above. This reporter cell line is a derivative of HeLa cells, but stably expresses high levels of CD4 and CCR5 on the cell surface, and has integrated copies of the luciferase gene under the control of the HIV-1 promoter that allows simple and quantitative analysis of HIV-1 infection. MT-4 cells were maintained in Roswell Park Memorial Institute 1640 (RPMI, Sigma, USA) medium supplemented with 10% fetal calf serum, 20 units/mL penicillin and 20 μg/mL streptomycin. Cells were transfected using FuGene HD according to the manufacturer’s instructions (Promega, USA) in a 3:1 ratio with DNA.

The pEGFPN1 and mCherry plasmids were obtained from Addgene (Addgene, USA). The plasmid encoding for HA-tagged ubiquitin was a kind gift from Soham Gupta (Karolinska Institutet, Sweden). pCR3.1/HIV-Gag-mCherry (HIV-1B Gag) and pCR3.1-GFP-ALIX plasmids were kind gifts from Dr. Paul Bieniasz (The Rockefeller University, USA). A codon-optimized HIV-1 Gag with PYQE motif gene of East African HIV-1C viruses was cloned into the pCR3.1/HIV-Gag-mCherry plasmid that contained an mCherry protein C-terminal tag sequence. The Gag-PYQE-mCherry plasmid was modified by site-directed mutagenesis to make a base pair substitution mutation (C to A) that changes the PYQE motif to PYKE using the Q5 SDM kit (New England Biolabs, USA). Gag-PYQE-mCherry plasmid was linearized by PCR using 5’-AAGGAGCCTCTGACGAGCC-3’ as forward primer (with the C to A base pair substitution underlined) and 5’-ATAGGGACCCTGGTCTTTCAGC-3’ as reverse primer. Linear products were re-circularized using DpnI, a polynucleotide kinase and a ligase from the SDM kit following the manufacturer’s protocol. DNA sequences were verified by Sanger sequencing.

### Gag-p6 sequences and clinical specimens

To identify the distribution of PYQE, PYRE and PYKE motifs, we re-analysed the HIV-1C gag-p6 sequences that were collected from three cohorts, the Swedish InfCare Cohort (n=140), the German Cohort (n=127), and the Ethiopian Cohort (n=73), as reported previously with respect to naturally occurring polymorphisms in PYxE motif (15). Additional HIV-1B (n=2754) and HIV-1C (n=1432) gag-p6 sequences were collected from HIV-1 Los Alamos Database. Stored plasma samples from therapy naïve patients infected with HIV-1C (n=10) were randomly selected based on gag-p6 sequences from the HIV-cohort at Karolinska University Hospital, Stockholm, Sweden.

### Recombinant virus production with patient-derived gag-pol

The recombinant viruses were produced as described by us recently (20). Briefly, the *gag-pol* fragment (HXB2:0702-5798) was cloned into pNL4-3 plasmid following digestion with *BssHII* and *SalI* (New England Biolab, USA) and ligation using T4 DNA ligase (New England Biolabs, USA). The chimeric viruses were produced by transient transfection of the plasmids into the 293T cell line using FuGene HD and harvested 72 hours later by a collection of the cell-free supernatant cleared by centrifugation and stored in aliquots at −80°C.

### In silico analysis: Molecular modeling, docking, and Ubiquitin binding motif prediction

Homology modeling techniques generated the structure of HIV-1C Gag late domain with the PYKE insertion. The published crystal structures of the Gag late domain in complex with ALIX (PDB entries 2XS1 (14) and 2R02 (21)) were used as template molecules to model HIV-1C Gag late domain. For this purpose, the ‘Prime’ utility of Schrödinger Suite (Schrödinger Inc., USA) was used. The structure was subjected to restricted minimization using OPLS_2005 force field. The resulting structure of HIV-1C Gag late domain structure was docked into the crystal structure of ALIX (PDB file 2XS1) using PIPER protein-protein docking program of BioLuminate Suite (Schrödinger Inc., USA), after deleting SIV_mac239_ PYKEVTEDL late domain structure. The resulting structure was further minimized for 1000 iterations to remove steric clashes. A similar protocol was used to form the complex among ALIX, Gag late domain, and ubiquitin. The crystal structure of ubiquitin in complex with the TSG101-binding PTAP domain (PDB file 1S1Q) was used after deleting TSG101 coordinates (22). The entire complex was then subjected to molecular dynamics simulations for 1,000,000 steps with step size of 50 picoseconds using an OPLS3 force field. The most energetically stable model of the complex was used for further analysis. To obtain energetically favored polar interactions, the conformational search of polar sidechains at the interface of proteins was also conducted.

The prediction of ubiquitin binding with the HIV-1C Gag late domain with the PYKE motif P6 was performed by adding the amino acid sequence in the prediction tools presented at the website http://www.ubpred.org/.

### Immunofluorescence

HEK293T cells were cultured on poly-L-lysine-coated glass coverslips and co-transfected with GFP-tagged ALIX and mCherry-tagged codon-optimized Gag variants. At 24 hours post-transfection, cells were washed twice with PBS, fixed in 10% formalin (Sigma, USA) for 20 min at room temperature, washed three times with PBS, and stored in PBS at 4°C. Later, nuclear counterstaining with DAPI was performed followed by mounting of the glass coverslips on glass slides. When HA-ubiquitin was also co-transfected, cells were fixed in ice-cold methanol for 15 min at −20°C, washed twice with PBS, and stored in PBS at 4°C. For ubiquitin detection, fixed cells were incubated with the anti-HA antibody (mouse monoclonal, clone 12CA5, Sigma, USA) for 1 hour at room temperature followed by goat-anti-mouse-Alexa Fluor 647-Plus-conjugated secondary antibody (Invitrogen, USA) for 1 hour at room temperature before nuclear counterstaining with DAPI. Fluorescence was analyzed by confocal laser scanning microscopy using a Nikon Single point scanning confocal microscope with ×60/1.4 oil objective (Nikon, Japan). Fluorescence intensity was measured along drawn lines using Fiji/ImageJ software (23). The highest fluorescence intensity value for each fluorophore along each line was set to 100%, and values were plotted as relative fluorescence intensity in percentage against distance in microns using GraphPad Prism v6 (Graphpad Inc., USA).

### Microscale Thermophoresis (MST)

The microscale thermophoresis experiments were conducted by Monolith NT0.115 instrument (NanoTemper Technologies) at 40% MST and LED power at 25°C. Hexahistidine-tagged ALIX protein was labeled by NTA-dye (NanoTemper Technologies) using manufacturer’s protocol. The peptides were synthesized at the Molecular Interaction Core (University of Missouri). Thermophoresis was induced by mixing peptides (0.001-1µM) and fixed NTA-labelled ALIX (50 nM) in a buffer 50 mM Tris-Cl pH 7.8, 100 mM NaCl and 0.1% pluronic-F127. The binding isotherms were obtained by plotting the difference in normalized fluorescence against increasing peptide concentration. The binding affinities were determined to fit the data points to a quadratic equation (equation 1) using non-linear regression using MO Affinity software (NanoTemper Technologies), Prism (GraphPad Inc. version 6.0) or OriginLab (version 18, OriginLab Corp. Northampton, MA, USA).

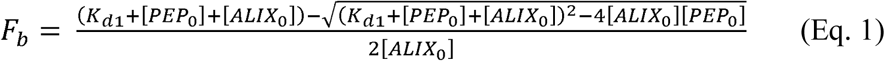

Where F_b_ is fraction of ALIX/peptide complex, where K_d_ = [ALIX][PEP]/[ALIX-PEP], [ALIX] is the concentration of free ALIX, [ALIX_0_] is the concentration of total ALIX, [PEP] is the concentration of free peptide and [PEP_0_] is the total concentration of peptide.

### Co-immunoprecipitation assay

HEK293T cells were co-transfected with GFP-tagged ALIX and codon-optimized Gag variants from HIV-1B or HIV-1C. At 24 hours post-transfection, cells were collected, washed twice with ice-cold PBS and immunoprecipitation of GFP-tagged proteins was performed using GFP-Trap®_A kit (Chromotek, Germany) according to the manufacturer’s protocol. Briefly, cells were lysed in ice-cold lysis buffer (10 mM Tris pH 7.5, 150 mM NaCl, 0.5 mM EDTA, 0.5% NP-40, supplemented with a protease cocktail inhibitor from Roche, Switzerland) for 30 minutes on ice. Then lysates were diluted in ice-cold wash buffer (10 mM Tris pH 7.5, 150 mM NaCl, 0.5 mM EDTA) supplemented with a protease cocktail inhibitor from Roche, input samples (10% of total) were saved for immunoblot analysis, and the remaining (90%) part of the diluted lysates were incubated with equilibrated GFP-Trap^®^_A beads end-over-end for 1 hour at 4°C. Beads were washed three times with ice-cold wash buffer before resuspending in 2x Leammli buffer (Invitrogen, 4x buffer diluted 1:1 with PBS) and heated for 10 minutes at 95°C. The protein concentration of input samples was determined by DC protein assay (Biorad, USA) according to the manufacturer’s microplate assay protocol. Equal amounts of protein for input samples and equal volumes of pull-down samples were subjected to immunoblotting using primary antibodies against GFP (rabbit monoclonal, clone EPR14104, Abcam, UK), Gag (rabbit polyclonal to HIV1 p55 + p24 + p17, Abcam, ab63917), and β-actin (rabbit polyclonal, Abcam, ab8227) and secondary horseradish peroxidase-conjugated antibodies (polyclonal goat anti-rabbit, DAKO/Agilent, USA). Immunoblotted proteins were detected using the enhanced chemiluminescence detection system (Pierce/Thermo Fisher Scientific, USA) and Hyperfilms (Amersham/GE Healthcare Life Sciences, UK).

### Viral growth kinetics assay in MT-4 cell

First, TCID_50_ for each propagated recombinant viral clone was determined in TZM-bl reporter cells. For each recombinant viral clone, ten-fold serial dilutions were prepared to range from 10^2^ to 10^7^ in dilution factor from a single aliquot in medium containing DEAE (20 µg/µL). TZM-bl cells were plated in 96-well plates, and six replicates were incubated with the various virus dilutions for 48 hours at 37°C in a 5% CO_2_ humidified incubator. Virus infectivity was quantified by measuring Renilla luciferase activity (relative light units [RLU]) using Bright-Glo™ Luciferase Assay System (Promega, USA) on the Tecan microplate reader (Tecan Infinite^®^ 200 Pro, Tecan Group Ltd, Switzerland). The Spearman-Karber method was used to calculate the TCID_50_ for each recombinant viral clone. Then, MT-4 cells (2 × 10^5^) were seeded in a 12-well plate and infected with each recombinant virus at a multiplicity of infection (MOI) of 0.05 plaque-forming units (pfu) per cell in triplicates for each condition. Supernatants were collected on day 0, 3, 5 and 7 days post infection, and HIV-1 p24 levels in supernatants were measured on the HIV COBAS 8000 platform (Roche Diagnostics, Switzerland). Viral growth kinetics (VGK) were analyzed as a relationship between signal to the cut-off ratio (i.e., electrochemiluminescence signal of the sample relative to calibrated negative samples for the HIV combi PT kit on the COBAS 8000 system) for each time point and replicate. Graphical representation of the result was made using Graphpad Prism v6 (Graphpad Inc., USA).

### Isolation of CD4^+^ T-cells

Primary CD4^+^ T-cells were isolated from donor peripheral blood mononuclear cells (PBMCs). Briefly, 4.3 × 10^8^ PBMCs were obtained from 50 ml buffy coat using the standard method of gradient separation using Ficoll Histopaque reagent followed by centrifugation. Isolated PBMCs were used to isolate CD4^+^ T-cells by negative selection using the EasySep ™ Human CD4^+^ T-cells isolation kit (Stem Cell Technologies, Canada). Briefly, PBMCs were pooled in 5 ml polystyrene round-bottom tubes at a concentration of 5 × 10^7^ cells and incubated with 50 µl of an antibody cocktail for 5 min at room temperature before mixing with RapidSphere ™ beads that was then followed by magnetic separation to yield pure CD4^+^ T-cells.

### Viral growth kinetics assay in CD4^+^ T-cells

CD4^+^ T-cells were cultured using RPMI media supplemented with 10% fetal calf serum and 1% pen-strep (penicillin and streptomycin). The cells were stimulated with PHA (20 µg/ml final concentration) for three days before starting the assay. To start the viral growth kinetics assay, 1 × 10^6^ CD4^+^ T-cells were seeded in each well in a V-bottom 96-well culture plate and infected with viruses at an M.O.I of 0.05 pfu/cell in the presence of 20 µg/ml of DEAE. The plate was spinoculated at 37°C for 2 hours at 800 rpm in a temperature-controlled centrifuge. After that, the plate was further incubated in a 5% CO_2_ incubator for 8 hours. Afterward, infected cells were transferred to a 48-well culture plate and washed six times before harvesting the initial supernatant from each corresponding well and designating it Day 0. The experiment was then monitored for seven days, and the supernatant was harvested for days 3, 5 and 7. HIV p24 was then measured in the supernatants using HIV alliance p24 ELISA kit (Pekin Elmer, USA) and results were analyzed using GraphPad Prism v6 (Graphpad Inc., USA).

### Drug sensitivity assay (DSA)

The following drugs were purchased from Selleckchem, USA: Atazanavir sulfate (ATV), darunavir ethanoate (DRV), lopinavir (LPV), azidothymidine (AZT), tenofovir alafenamide (TAF), efavirenz (EFV), rilpivirine (RPV), etravirine (ETR), raltegravir (RAL), elvitegravir (EVG), dolutegravir (DTG), and cabotegravir (CAB).. The DSA was performed by determining the extent to which the antiretroviral drug inhibited the replication of the reference virus (pNL4-3), wild-type, PTAPP duplicated and PYxE derived recombinant viruses, respectively. One round was required for all drugs as previously reported (20), except for the protease inhibitor (PI) drugs that required two rounds of infection. Briefly for the for two-round infection using PI DSA, TZM-bl cells (1×10^4^) were seeded in 96-well plates and cultured for 24 hours. Drugs were serially diluted in culture media (ranging from 0.1 µM to 0.01 pM) and added in triplicate to the cells. The following day, viruses were added to each well at an MOI of 0.05 pfu/cell in the presence of DEAE (10 µg/ml). After 48 hours, the viral supernatant from respective 96-wells plates with each drug was transferred to newly TZM-bl pre-seeded 96-wells plates without adding new drugs. After that, virus replication was quantified by measuring Renilla luciferase activity (relative light units [RLU]) using Bright-Glo™ Luciferase Assay System (Promega, USA) 48 hours post-reinfection. Drug concentrations required for inhibiting virus replication by 50% (EC_50_) were calculated by a dose-response curve using non-linear regression analysis (GraphPad Prism, version 6.07; GraphPad Software, USA). The DSA experiments were performed with three technical replicates for each virus with the specified dynamic concentration range of the drug, and at least two independent analyses were performed. The reproducibility of the DSA was assessed by the 95% confidence interval obtained for the drug EC_50_ and the degree of correlation between technical replicates. The output for the drug EC_50_ results were used to compute the fold change value (FCV) for each virus relatively to pNL4-3.

## Results

### Evolution of gag-p6 late domain sequences in HIV-1B and HIV-1C

The sequence analysis of HIV-1B and South African and Indian HIV-1C gag-p6 sequences (obtained from the Los Alamos Database) identified a conserved LYPx_n_L motif in HIV-1B which was missing in HIV-1C (Fig 1a). We noted a unique evolution of the lysine (K) residue in the HIV-1B and HIV-1C gag-p6 late domain, which is the target of ubiquitination. All the amino acids positions are per the standard HXB2 co-ordinates. While K475 residue was conserved in both HIV-1B and C, there was an evolution of a lysine residue at amino acid position 479 along with K481R mutation in the majority of HIV-1C sequences. Interestingly, the multiple sequence alignment of HIV-1C_PYxEi_ strains identified conservation of HIV-1C specific K479 residue, but residue R481 in HIV-1C_WT_ changed to a lysine residue within HIV-1CPYxEi strains and thus identical to K481 in HIV-1B (Fig 1a). When we analyzed gag-p6 sequences from HIV-1C patients from the Swedish InfCare HIV cohort (n=140), we found that 68% (95/140) of HIV-1C patients had a wild type strain whereas 32% (45/140) had a PYxE insertion; 22% (31/140) had a PYKE insertion, and 9% (13/140) had a PYQE insertion (Fig. 1b). All of the PYxE sequences in HIV-1C of patients from the Swedish cohort belonged to HIV-1C strains of Ethiopian and Eritrean origin. Similar trend was observed in the German HIV-1C cohort with 12% PYKE insertion (15/127) (Fig. 1c), and in the Ethiopian HIV-1C cohort with 37% (27/73) PYKE insertion (Fig. 1d). Thus, our analysis indicated that PYKE and PYQE are predominating in HIV-1C_PYxEi_ viruses of all cohorts analysed. Subsequently, we restricted our analysis to these two variants.

**Figure 1.**
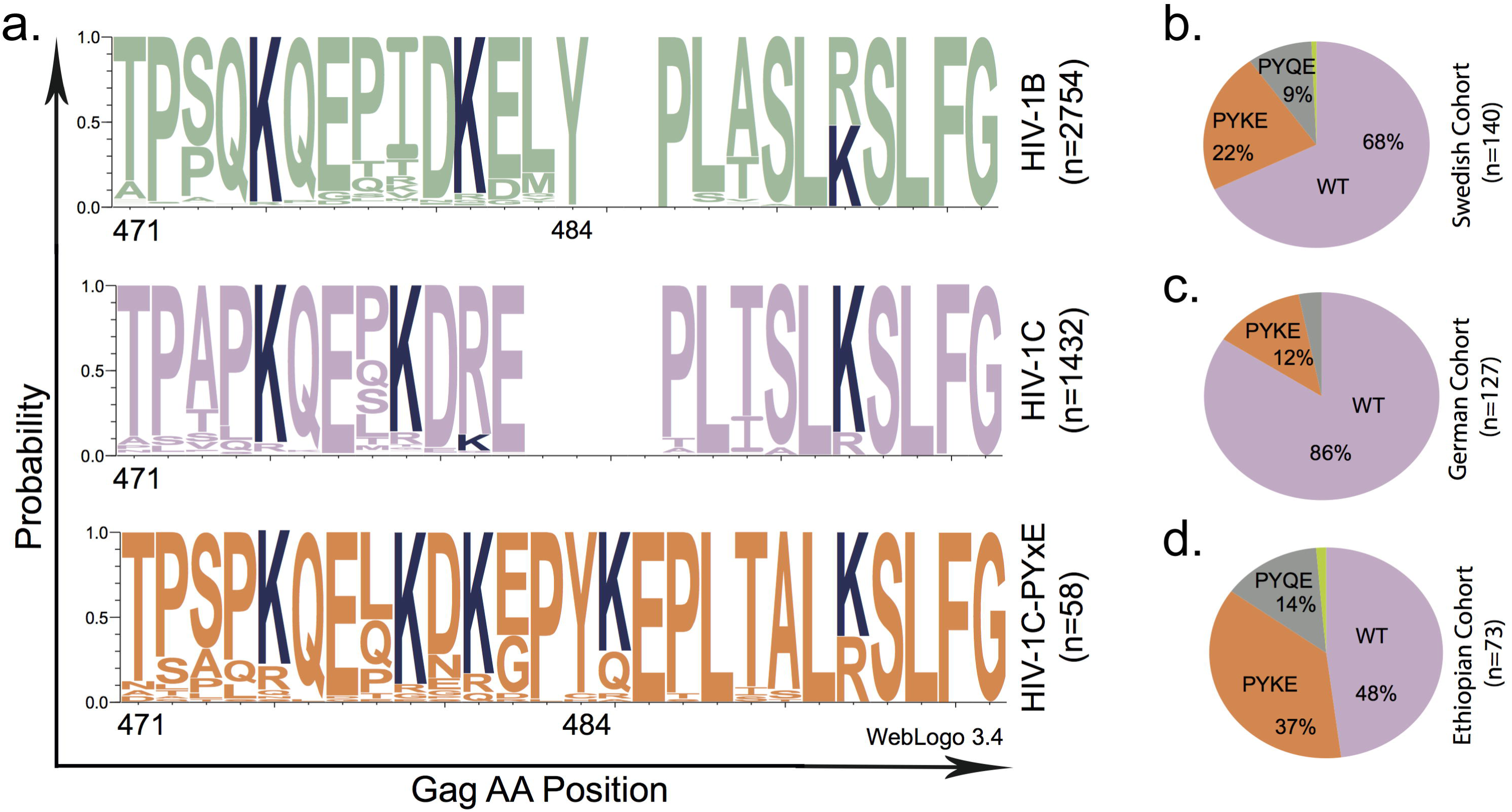
Consensus sequences and distribution of HIV-1 gag. **(a)** Aligned consensus sequences of a gag from HIV-1 type B wild-type (top), HIV-1 type C wild-type (middle), and HIV-1 type C with tetrapeptide insertion (bottom) from the Los Alamos database. Sequence logos with one-letter coded amino acid were generated using WebLogo 3. **(b-d)** Distribution of HIV-1 type C with gag_wt_, gag_PYKE_, and gag_PYQE_ in patient cohorts from Sweden **(b)**, Germany **(c)** and Ethiopia **(d)**.

### PYxE insertion enhances type C Gag: ALIX interaction

Prior to testing the interaction between Gag and ALIX, we evaluated the cellular localization of both proteins. 293T cells were co-transfected with GFP-tagged ALIX with or without three codon-optimized variants of mCherry-tagged HIV-1C Gag, namely wild type Gag or Gag that includes the tetrapeptide insertion PYKE or PYQE. Confocal analysis showed cytoplasmic localization of ALIX and predominant plasma membrane localization of all three Gag variants (HIV-1C_WT_, HIV-1C_PYQEi_ and HIV-1C_PYKEi_) as the complete Gag produces the virus-like particles (Fig. 2). Thus, the tetrapeptide insertion did not cause a change or defect in the localization pattern of Gag proteins at the plasma membrane.

**Figure 2.**
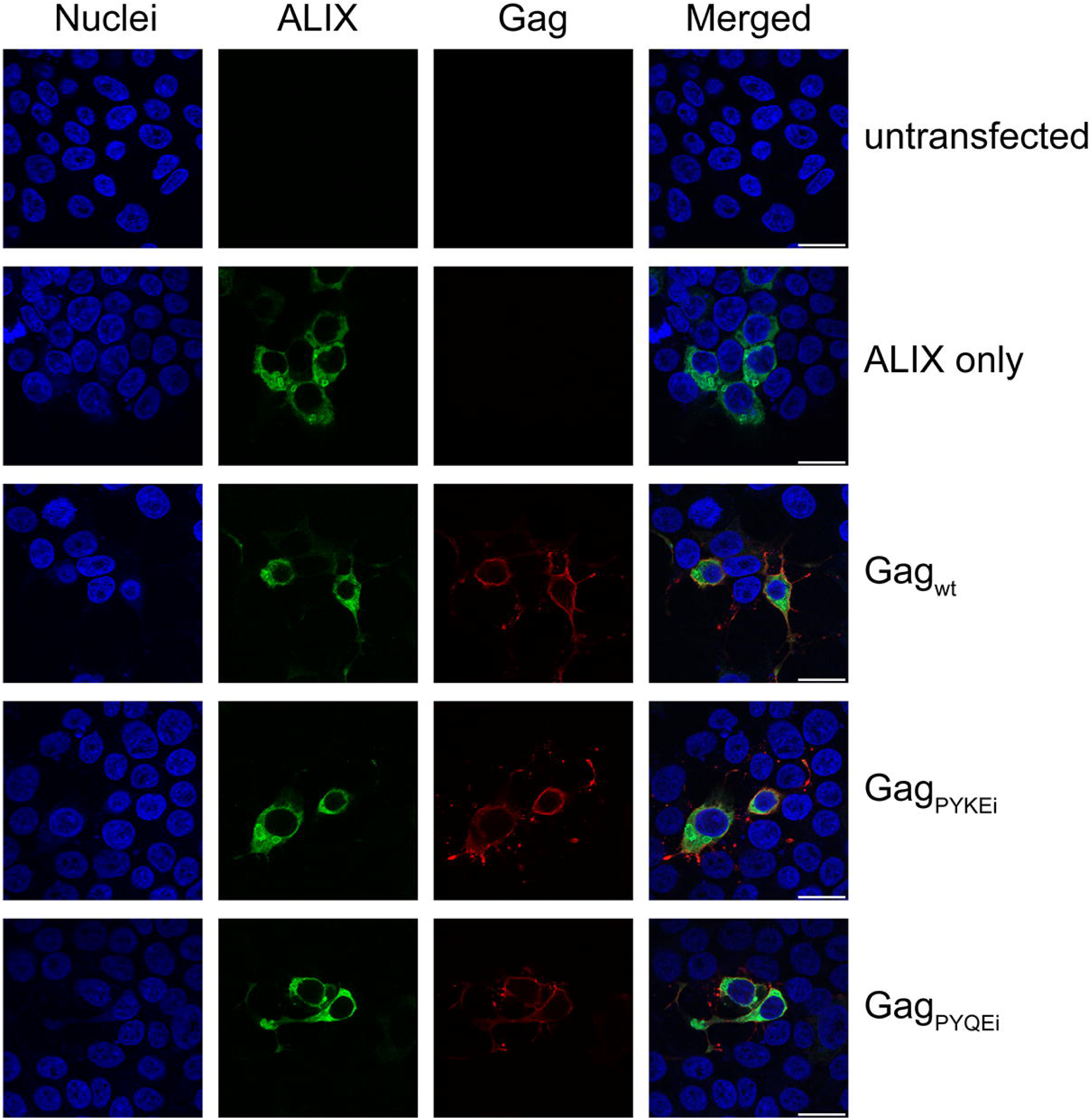
Cellular localization of Gag variants and ALIX. Confocal images of HEK293T cells co-transfected with the GFP-tagged ALIX and mCherry-tagged HIV-1C Gag variants (wt, PYKEi, & PYQEi) expression plasmids. Nuclei are visualized in blue, ALIX in green, and Gag in red. Bars, 20 μm.

Next, we evaluated the binding affinity of four HIV-1 Gag variants, HIV-1B_WT_, HIV-1C_WT_, HIV-1C_PYKEi_ and HIV-1C_PYQEi_, to ALIX (Fig 3a). The late domain motifs’ peptides were selected based on crystal structure PDB:2XS1 and PDB:2R02. The binding affinity of the Gag peptide with ALIX were statistically significantly higher in HIV-1B_WT_ (32 ± 3 nM) compared to all HIV-1C variants; HIV-1C_WT_ (290 ± 6 nM), HIV-1C_PYKEi_ (110 ± 4 nM), HIV-1C_PYQEi_ (96 ± 4 nM) (Fig 3b). However statistically significant higher binding affinity to ALIX was observed with HIV-1C_PYKEi_ and HIV-1C_PYQEi_ compared to HIV-1C_WT_.

**Figure 3.**
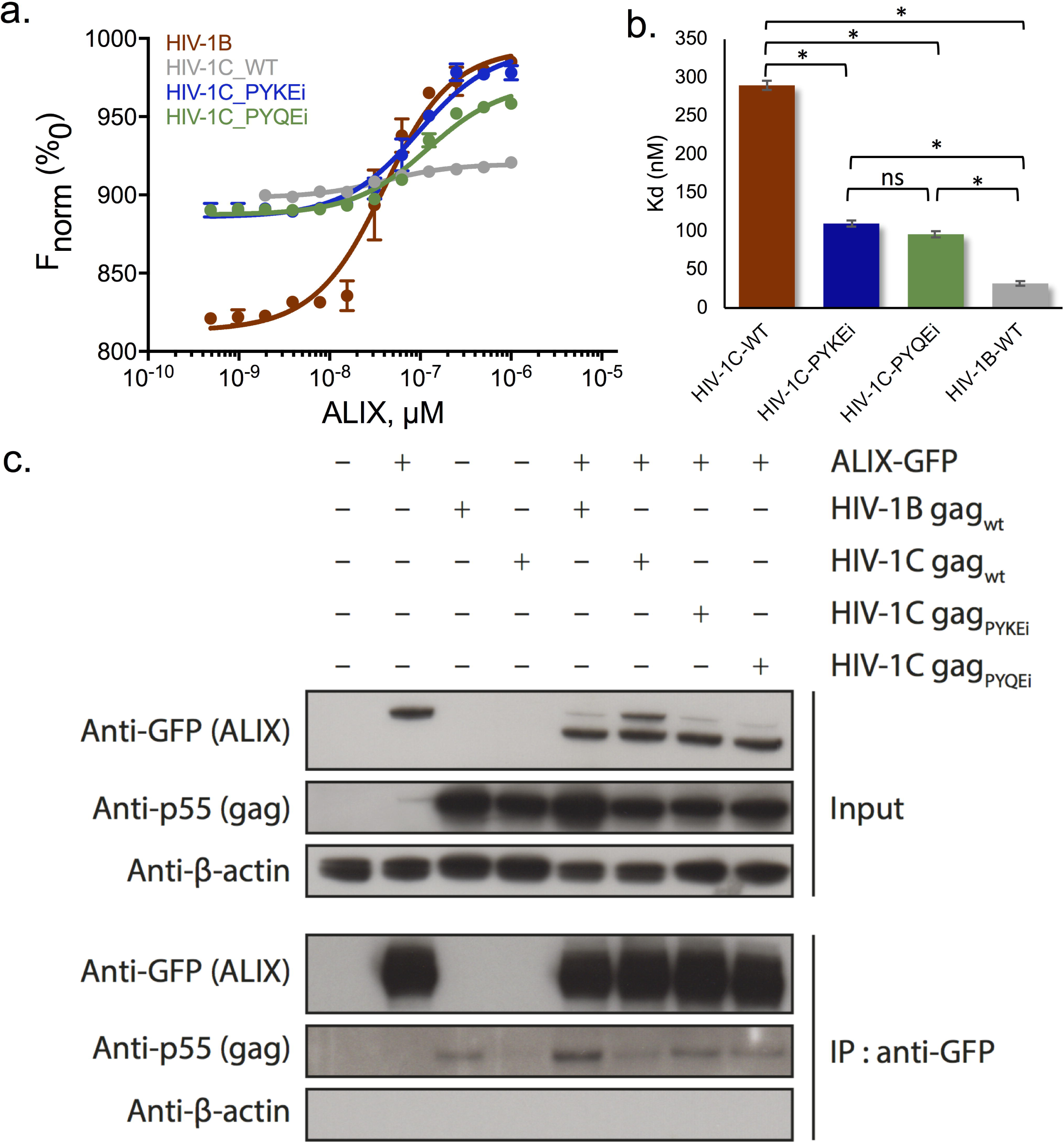
HIV-1C Gag with PYxE insertion has increased binding with ALIX. **(a-d)** Protein-peptide interaction using Microscale thermophoresis. The peptide sequences are given in the individual graphs. **(e)** The equilibrium dissociation constant (Kd) of the individual interactions. The smaller the Kd value, the greater the binding affinity of the ligand for its target. **(f)** HEK293T cells were co-transfected with GFP-tagged ALIX and Gag variants from HIV-1 type B or C. At 24 hours post-transfection, cells were lysed and subjected to GFP-pull down. Input and pull-down samples were subjected to immunoblotting antibodies against GFP (ALIX), p55 (Gag), and β-actin. Data depicted are representative of at least two independent experiments.

As the peptides may not retain their conformational structures as in the complete protein, we confirmed the binding by co-immunoprecipitation pull-down assays. GFP-tagged ALIX and HIV-1C Gag variants were transfected either alone or together in 293T cells. A plasmid encoding for HIV-1B Gag was used together with GFP-ALIX as a positive control for Gag:ALIX interaction. Lysates were prepared 24 hours post-transfection, GFP-tagged proteins were pulled down and co-immunoprecipitated proteins were analysed by immunoblotting (Fig. 3c). Pull down of GFP-tagged ALIX was efficiently and the amount of HIV-1B Gag was increased when it was co-transfected with GFP-ALIX confirming that HIV-1B Gag binds to ALIX. HIV-1C Gag wild type was also detected when co-transfected with GFP-ALIX, but HIV-1C Gag PYKE was more prominently detected. The intensity of HIV-1C Gag PYQE was consistently lower than that of the PYKE variant, but slightly increased compared to the wild type HIV-1C Gag. Thus, the tetrapeptide insertion PYxE in HIV-1C Gag-p6 late domain promotes binding of Gag towards ALIX.

### In silico analysis predicts binding of ALIX to the PYKE motif in Gag facilitated by ubiquitination

As reported before, the lysine residues in the Gag-p6 are subjected to ubiquination (24), we first predicted the likelihood of ubiquitination of the lysine residue of PYKE (K_ins_) compared to lysine residues adjacent to the PYKE motif. Indeed, K_ins_ was predicted to be more likely ubiquitinated than lysine residues in the vicinity of the PYKE motif (Fig. 4a). Since the PYKE motif is present at the missing ALIX-binding site and contains three charged amino acids that can facilitate interaction with other proteins, we used the PYKE insertion (amino acids 483 to 486) in the molecular modelling to predict the interaction of the Gag-p6 late domain with ALIX. Amino acids E482, Y484 and E486 of Gag could directly facilitate binding to ALIX (Fig.4b and Table 1). Ubiquitin was predicted to bind K501 in ALIX and thereby affected the interaction between E486 in Gag-PYKE with K501 in ALIX. In addition, ubiquitin could also interact with E387 of ALIX and K_ins_ of Gag-PYKE through its residues K63 and E64, respectively. Thus, our *in silico* analysis predicted that the PYxE motif can restore the interaction of Gag with host cell ESCRT-III adaptor protein ALIX and that, in case of a PYKE insertion, ubiquitination of the K_ins_ residue could further facilitate this interaction. We also tested the localization of ubiquitin by immunofluoresence. Since both Gag and ALIX are known to be ubiquitinated, it is not surprising that ubiquitin co-localized with ALIX and Gag (Fig. 4c).

**Figure 4.**
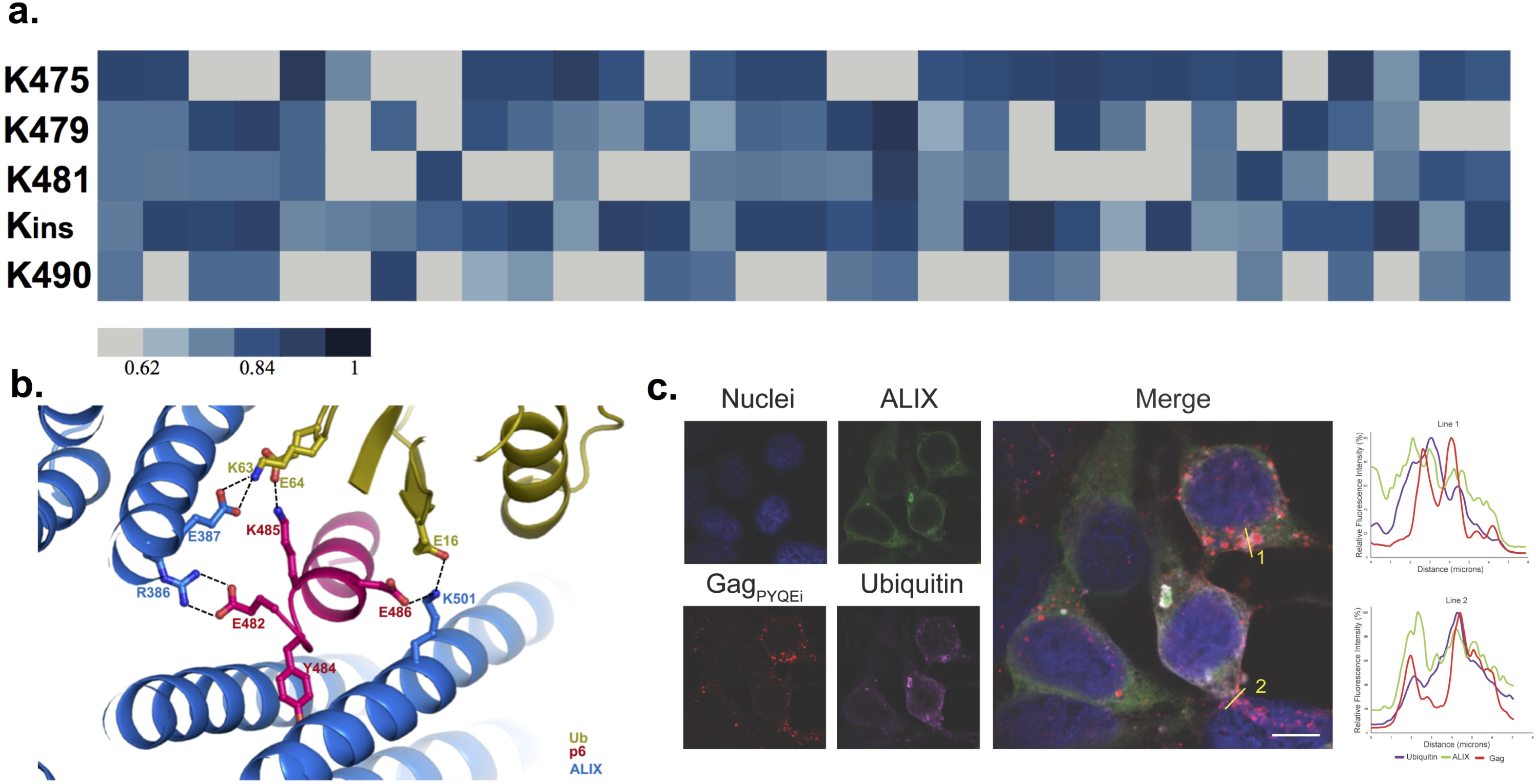
*In silico* binding prediction between PYKE inserted within HIV-1 type C Gag, ALIX, and ubiquitin. **(a)** Likelihood of ubiquitin binding to lysine residues in the vicinity of or within the PYKE motif. K_ins_ represents the lysine residue within the PYKE motif. **(b)** All proteins are presented as ribbon-like structures. The red ribbon represents Gag and its PYKE insertion, the blue ribbon ALIX and its Gag-binding sites, and the green ribbon ubiquitin. Amino acids involved in Gag: ALIX binding is shown in one-letter format. Interactions between amino acids are shown with black dotted lines. *In silico* docking was performed using Schrödinger software. **(c)** Confocal images of HEK293T cells transfected with the GFP-tagged ALIX and mCherry-tagged HIV-1C Gag PYQEi and HA-tagged ubiquitin expression plasmids. Nuclei are visualized in blue, ALIX in green, Gag in red, and ubiquitin in magenta. Bar, 20 μm.

**Table 1.** List of amino acids that are involved in the interaction between Gag-P6, ALIX and ubiquitin (Ub).

| Interacting atoms | Residue 1 | Residue 2 | Distance (Å) | Interaction type |
| --- | --- | --- | --- | --- |
| OE1 – NH1 | E482 (LD) | R386 (ALIX) | 2.96 | Charge-charge |
| OE2 – NH2 | E482 (LD) | R386 (ALIX) | 2.83 | Charge-charge |
| OE1 – NZ | E387 (ALIX) | K63 (Ub) | 3.05 | Charge-charge |
| OE2 – NZ | E387 (ALIX) | K63 (Ub) | 2.97 | Charge-charge |
| OE2 – NZ | E64 (Ub) | K485 (LD) | 2.93 | Charge-charge |
| OE2 – NZ | E486 (LD) | K501 (ALIX) | 2.95 | Charge-charge |
| OE1 – NZ | E16 (Ub) | K501 (ALIX) | 3.12 | Charge-charge |
| OH – OD1 | Y484 (LD) | D506 (ALIX) | 2.72 | Hydrogen Bond |

E = glutamic acid, R = arginine, Y = tyrosine, D = aspartic acid, K = lysine

### Differences in viral growth of clinical strains of HIV-1C with PYxE motif

Finally, we assessed the consequence of the PYxE insertion on HIV-1C replication fitness using clinical strains obtained from HIV-1C infected patients. Gag-pol sequences were cloned from one wild type HIV-1C virus with a single PTAP motif and no PYxE insertion (PT01), three HIV-1C strains with a PYKE insertion (PT04-06), and one with a PYQE insertion (PT07). We also included two HIV-1C strains with a PTAP duplication (PT02 and PT03) as it has been shown to increase the viral fitness and affect the drug sensitivity (Fig. 5a) (25, 26). Then, these sequences were cloned into the genome of a reference virus (NL4-3, HIV-1B wild type) replacing its gag-pol sequence. The various recombinant viruses were tested for viral replication in an *ex vivo* viral growth kinetic assay and compared to the reference virus in MT-4 cell line (Fig. 5b) and primary CD4^+^ T-cells isolated from donor blood (Fig 5c). In MT-4 cells, a statistically lower amount of HIV-1C_WT_ virus was observed in the supernatant compared to the HIV-1C_PYxEi_ and HIV-1B reference viruses three days post infection. At day five post infection, the amounts of HIV-1C_WT_ virus was still reduced compared to the other viruses and reached the highest values at day 7 post infection. The reduction of HIV-1C_PYxEi_ and HIV-1B viruses at day 7 is most likely caused by the dead of the virus-producing cells, which was observed microscopically. Similar data was observed in CD4^+^ T-cells where HIV-1C_WT_ showed the lowest amount of virus in the supernatant compared to all other viruses at day 7 (Fig. 5c). Our data that viral replication for the HIV-1C wild type clone is reduced compared to the HIV-1B reference virus is in concordance with a previous report (27). More importantly, our data showed that when a PTAP-duplication or a PYxE insertion was present, viral growth was increased compared to the HIV-1C wild type variant and was comparable to HIV-1B wild type. This correlation suggests that HIV-1C strains with a PYxE insertion have a growth advantage compared to wild type HIV-1C. This is also in line with our earlier study that HIV-1C_PYQEi_ viruses are more replication competent as well as more pathogenic (17).

**Figure 5.**
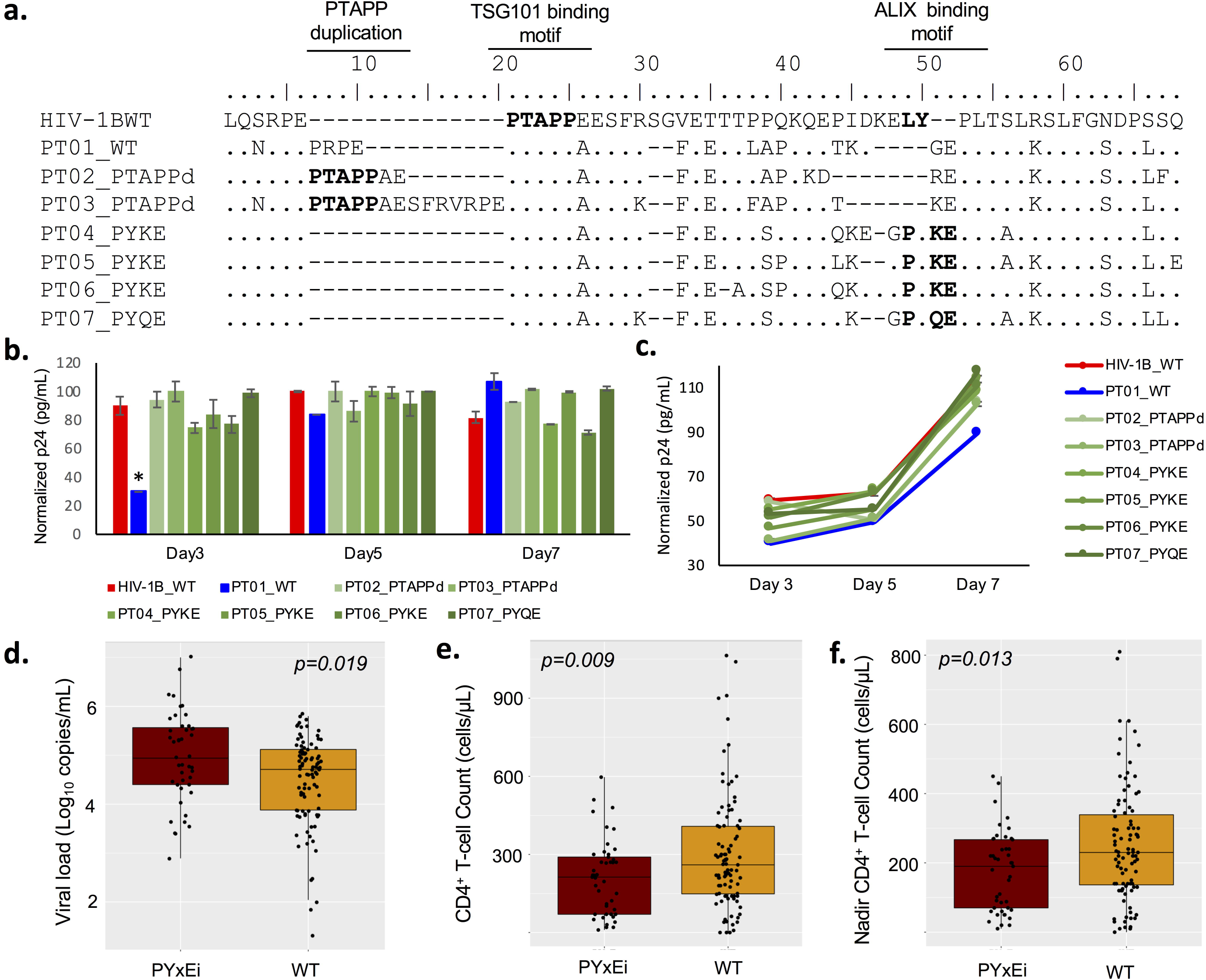
PYxE insertion increases viral growth and drug sensitivity towards protease inhibitor drug lopinavir. **(a)** Gag-pol sequences were amplified from HIV-1 type C patient isolates and cloned into HIV-1 molecular clone pNL4-3. Recombinant viruses were propagated in HEK293T cells and MOIs were determined in TZM-bl reporter cells. Viral growth assay was performed in MT-4 cells **(b)** or purified primary CD4+ T cells **(c)**. Cells were infected with the various recombinant viruses at an MOI of 0.05 and supernatants were harvested on day 0, 3, 5, and 7. Viral load in supernatants were measured by HIV-1 p24 analysis on the HIV COBAS 8000 platform. Data represent the mean ± SD of triplicates. Clinical data regarding viral load at initiation of therapy **(d)**, CD4+ T cell count at initiation of therapy **(e)**, and nadir CD4+ T cell count **(f)** was collected from the Swedish HIV-1C cohort and patients were grouped as HIV-1C_PYxEi_ (n=45) or HIV-1C_WT_ infected individuals (n=95).

Next, we checked whether the *ex vivo* data corroborate with the clinical data. In the Swedish HIV-1C cohort (n=140) we checked the nadir CD4^+^ T-cell count, CD4^+^ T-cell count at the initiation of therapy, and viral load at initiation of therapy. The viral load was significantly higher in HIV-1C_PYxEi_ infected individuals compared to the individuals who were infected with wild type HIV-1C viruses (Fig 5d). The CD4^+^ T-cell count at the initiation of therapy (Fig 5e) and nadir CD4^+^ T-cell count (Fig 5f) were also statistically lower in the HIV-1C_PYxEi_ infected individuals compared to individuals infected with wild type HIV-1C viruses. These data further strengthens our *ex vivo* findings.

### Effect of PYxE strains in susceptibility towards antiretroviral drugs

Several reports have shown that a PYxE insertion is more frequent among HIV-1C therapy failure patients (16, 28). To examine whether our various recombinant viral clones without any known drug resistance mutations to RTIs, PIs and INSTIs responded differently to various ART drugs, the target cells were infected with individual viral clones in the presence of a specific drug. The viral replication was assessed and the EC_50_ was calculated and fold change of the EC_50_ was mentioned (Fig. 6). No differences in EC_50_ values were observed among the various recombinant viral clones in the presence of non-nucleoside RTIs (NNRTIs) efavirenz (EFV), rilpivirine (RPV) or etravirine (ETR), or INSTIs raltegravir (RAL), elvitegravir (EVG), dolutegravir (DTG) or cabotegravir (CAB). In contrast, with the PI lopinavir (LPV) one out of two viral clones with a PTAP-duplication and two out of four with PYxE insertion showed a >4-fold decreased sensitivity towards the drug. However, there was no change in susceptibility against DRV and ATV. One of the other viral clones with a PYxE insertion showed a 3.5-fold decreased sensitivity towards nucleoside reverse transcriptase inhibitor (NRTI) tenofovir alafenamide (TAF). Thus, our data indicate that PYxE insertion may potentially alter the sensitivity towards the PI LPV and NRTI TAF and that may be due to the increased replication fitness. However, this is not universal across all the HIV-1_PYxEi_-strains and could be due to co-evolution of other amino acids in the Gag protein.

**Figure 6.**
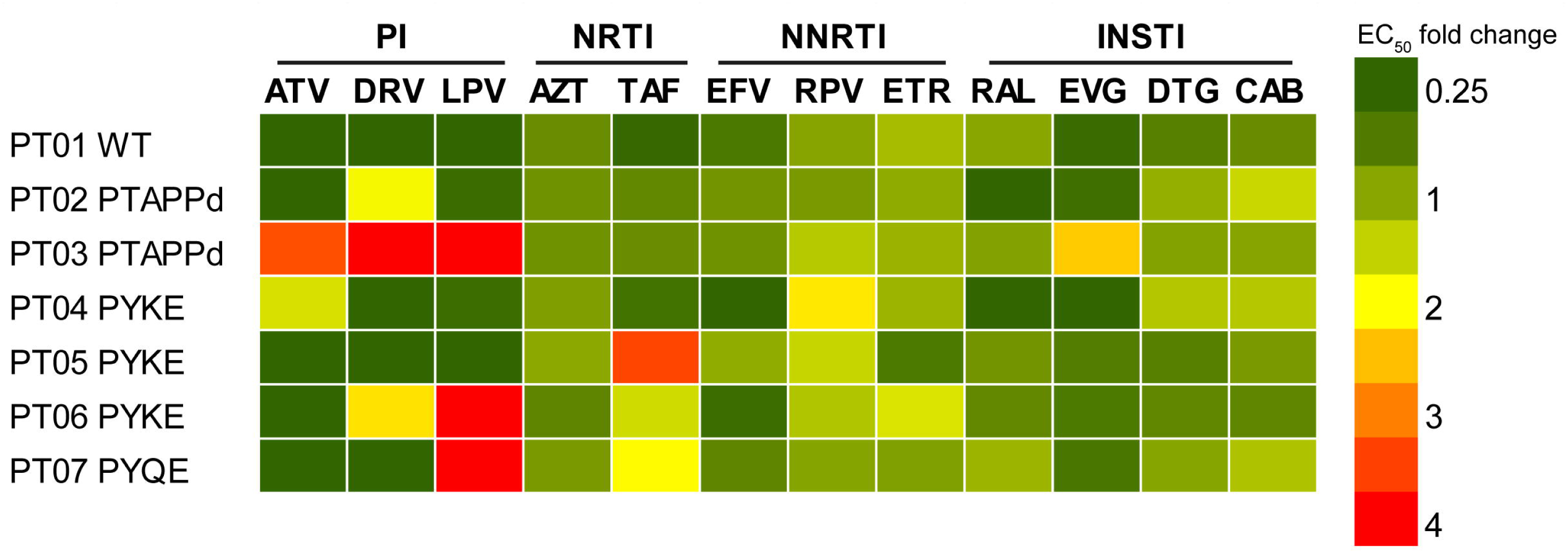
PYxE insertion increases drug sensitivity towards protease inhibitor drug lopinavir. TZM-bl reporter cells were infected with the individual recombinant HIV viruses at an MOI of 0.05 and cultured in the presence of individual anti-retroviral drugs at different dilutions. EC_50_ values for each virus with each drug were determined and EC_50_ fold changes (FC) were calculated compared to pNL4-3 reference virus. PI, protease inhibitor; NRTI, Nucleoside Reverse Transcriptase Inhibitor; NNRTI, Non-Nucleoside Reverse Transcriptase Inhibitor; INSTI, Integrase Strand Transfer Inhibitor. ATV, Atazanavir sulfate; DRV, darunavir ethanoate; LPV, lopinavir; AZT, azidothymidine; TAF, tenofovir alafenamide; EFV, efavirenz; RPV, rilpivirine; ETR, etravirine; RAL, raltegravir; EVG, elvitegravir; DTG, dolutegravir; CAB, cabotegravir. The FC data presented in the heat map was the mean of three individual experiments.

## Discussion

Genetic diversity among HIV-1 strains and their adaptation can provide advantages for individual viral strains in different biological contexts. Here, we described the distribution of HIV-1C containing tetrapeptide PYxE insertion within Gag and assessed the binding efficiency of Gag with host cell protein ALIX. Our results showed that the PYxE insertion enhances the binding capacity of HIV-1C Gag to ALIX in a potentially ubiquitin-dependent or independent manner. Also, the insertion increased the replication fitness of the virus in both an *in vitro* model T cell line as well as in primary CD4^+^ T cells and seemed to alter the sensitivity against the PI LPV and NRTI TAF in some of the strains.

Although the PYxE insertion is quite prevalent among HIV-1C from Ethiopia and Eritrea individuals, it is not as prevalent in South African or Indian HIV-1C strains (16, 28). The twelve base pair sequence encoding the PYxE motif is not present in non-subtype C M group HIV-1 sequences but is present within the Gag-p6 late domains of SIV_mac239_, SIV_smE543_, and their close relative HIV-2 (14, 29). As HIV-1C has a natural deletion of the YPx_n_L motif and insertion of PYxE has replication advantages, we hypothesize that these strains have evolved through a recombination event of HIV-1C with either SIV or HIV-2. It is well described that the HIV-1 epidemic in Ethiopia geographically clusters very strongly (30) and the Ethiopian HIV-1C has been proposed to originate from either a single lineage or multiple descendants (31). However to explain why the PYxE insertion is significantly more prevalent in HIV-1C from Ethiopia and Eritrea as compared to South African and Indian strains needs a further evolutionary study to prove this hypothesis.

The Gag-p6 late domain mediates viral budding through the PTAP motif, the YPx_n_L motif, and the PYxE motif. The PTAP motif mediates binding of Gag to ESCRT protein TSG101 and is present in all HIV and SIV variants. Therefore, this domain appears to be necessary for viral budding. Although one report shows that PTAP duplication only increases HIV-1 replication in the presence of a PI (26), others and we observed increased viral growth when HIV-1 contained a PTAP duplication (25). Although deletion of the PTAP motif has not been identified in clinical strains so far, artificial deletion or mutation of this PTAP motif severely abrogates release of HIV-1 from infected cells (11, 32, 33). However, the presence of either one of the two other motifs that both facilitate binding to host cell protein ALIX still allows for virion release (12, 14, 19, 34, 35). This indicates that both TSG101- and ALIX-mediated pathways for viral budding are not dependent on each other and each pathway is sufficient to mediate viral budding. The first ALIX-binding motif is YPx_n_L and is present in most HIV-1 subtypes but not in HIV-1C or HIV-2 (13, 14). HIV-2 Gag can mediate binding to ALIX through a distinct PYKEVTEDL motif, and mutations in this motif can abrogate ALIX binding and decrease virion release (14, 29). Thus, the PYxE motif in a subgroup of HIV-1C strains is similar but shorter than the ALIX-binding motif found in HIV-2. We showed that this insertion of PYxE in HIV-1C Gag at the natural deletion site of the YPx_n_L domain could reconstitute binding of HIV-1C Gag to ALIX. Hence, together with our observed PYxE insertion in HIV-1C, all HIV-1 subtypes have an ALIX-binding motif next to their PTAP-motif within the Gag late domain.

Our data suggest that ubiquitin may play a role in facilitating the binding of ALIX to the PYKE motif. This is not surprising, as ubiquitin has been shown to be involved in viral budding of HIV. Although dispensable for virus budding, ubiquitination of Gag and the ESCRT-proteins are essential for efficient release of HIV-1 from infected cells (24, 36, 37). Ubiquitination of the lysine of the PYKE motif could increase the binding between ALIX and Gag and thereby mediate more efficient viral budding. Indeed, co-immunoprecipitation of HIV-1C Gag-PYKE was more efficient compared to the Gag-PYQE variant. As the PYKE motif insertion seems to be the original insertion within HIV-1C Gag, ubiquitination of its lysine might not be essential since other variants of the PYKE motif, i.e., PYQE and PYRE, are identified among HIV-1C strains along with the evolution of another lysine (K) residues in the late domain. Because we only have few clinical strains with a PYxE insertion, we could not identify differences in viral growth between HIV-1C Gag-PYKE and other PYxE variants. Nonetheless, whereas the other three residues seem to be required for the binding of ALIX to Gag, the lysine residue might provide increased stability for this interaction.

Although the ALIX-mediated viral budding mechanism appeared to be lost in HIV-1C, it is surprising to observe that HIV-1C PYxE strains were prevalent among HIV-1C infected individuals. This suggests that the interaction of Gag with ALIX is of biological importance for the virus. In our limited cohort of HIV-1C infected patients, PYxE insertion was always co-present with the PTAP motif. This suggests that the PYxE motif is not redundant from the PTAP motif-mediated viral budding process through TSG101 nor a compensatory mechanism for loss of the PTAP-motif and HIV-1 requires both motifs for efficient viral replication. Importantly, viruses with HIV-1C Gag PYKE or PYQE insertion showed enhanced viral growth compared to viruses with HIV-1C wild type Gag (Fig. 5). This finding is in line with a recent report that showed that a PYRE insertion within Gag enhances HIV-1C viral growth (19).

Decreased sensitivity towards PI LPV or NRTI TAF was found in several HIV-1C strains with a PTAP duplication or PYxE insertion. For the PTAP duplication, it has previously been reported that it could enhance HIV-1 growth in the presence of PIs lopinavir and ritonavir (26). For the PYxE insertion, our earlier studies have shown that the PYxE insertion is more frequent in HIV-1C therapy-failure patients (15, 16). Nonetheless, since our clinical HIV-1C PYxE strains did not have any known drug resistance mutations, it seems likely that the correlation of these viruses with advanced viral growth and reduced sensitivity towards the PI LPV and TAF could increase the risk for therapy failure in HIV-1C PYxE-infected patients. Mechanistic studies show that at the time of reverse transcription of HIV-1C from the RNA template, the RT favors pausing at the nucleotides in HIV-1C K65 position (AAG) and this correlates with increased probability for the development of TDF/TAF mutation K65R (38, 39). However, no studies have shown any co-relation with Gag-mutation on the susceptibility of TAF. We are presently performing research to understand the role of Gag mutations on sensitivity for TAF as both tenofovir (TDF) and TAF are used globally.

In conclusion, our study showed that PYxE insertion in HIV-1C of patients originating from Ethiopia and Eritrea restored the interaction of Gag with ALIX, which is mediated in an ubiquitin-dependent or -independent way. Importantly, the insertion was positively correlated with the replication fitness that could affect the sensitivity against LPV in the absence of any PI drug resistance mutation. Based on the present study, our earlier clinical studies (15-17) and study by Chaturbhuj et al. (19), we, therefore, posit that PYxE insertion in HIV-1 subtype C strains provide replication advantage which can affect the susceptibility to certain antiretroviral drugs. As PYxE insertion evolved in treatment failure patients from HIV-1C from India and South Africa, it also provides replication advantage following the treatment failure and evolution of drug resistance mutations that may cost replication fitness. Most importantly, reduced sensitivity against TAF in the absence of any TAF-mutation needs further studies to analyze any role of Gag-mutations in reducing susceptibility towards TAF. Altogether, it will be important to follow up on whether HIV-1C_PYxEi_ strains are emerging following the failure of antiretroviral therapy with LPV and TAF based regimens due to increased replication fitness. This information could provide important insights in the clinical significance of the PYxE insertion within HIV-1C Gag.

## Acknowledgments

The study is funded by grants from the Swedish Research Council (2017-01330, UN; 2016-01675, AS), Karolinska Institutet Doctoral Student Funding (KID2015-154, UN) and the Stockholm County Council (ALF 20160074, AS). We thank S.G. Aralaguppe, W. Zhang and S. Svensson Akusjärvi for technical assistance. The microscopic part of the study was performed at the Live Cell Imaging facility, Karolinska Institutet, Sweden, supported by grants from the Knut and Alice Wallenberg Foundation, the Swedish Research Council, the Centre for Innovative Medicine and the Jonasson center at the Royal Institute of Technology, Sweden. The following reagents were obtained through the AIDS Reagent Program, Division of AIDS, NIAID, NIH: TZM-bl cells (Cat#8129) from Dr. John C. Kappes, and Dr. Xiaoyun Wu, MT-4 cells from Dr. Douglas Richman, and HIV-1 NL4-3 Infectious Molecular Clone (pNL4-3) from Dr. Malcolm Martin (Cat#114).

## References

1. Siddik AB, Haas A, Rahman MS, Aralaguppe SG, Amogne W, Bader J, Klimkait T, Neogi U. 2018. Phenotypic co-receptor tropism and Maraviroc sensitivity in HIV-1 subtype C from East Africa. Sci Rep 8:2363.

2. Neogi U, Haggblom A, Santacatterina M, Bratt G, Gisslen M, Albert J, Sonnerborg A. 2014. Temporal trends in the Swedish HIV-1 epidemic: increase in non-B subtypes and recombinant forms over three decades. PLoS One 9:e99390.

3. Sundquist WI, Krausslich HG. 2012. HIV-1 assembly, budding, and maturation. Cold Spring Harb Perspect Med 2:a006924.

4. Bieniasz PD. 2006. Late budding domains and host proteins in enveloped virus release. Virology 344:55–63.

5. Demirov DG, Freed EO. 2004. Retrovirus budding. Virus Res 106:87–102.

6. Votteler J, Sundquist WI. 2013. Virus budding and the ESCRT pathway. Cell Host Microbe 14:232–241.

7. Martin-Serrano J, Neil SJ. 2011. Host factors involved in retroviral budding and release. Nat Rev Microbiol 9:519–531.

8. Scourfield EJ, Martin-Serrano J. 2017. Growing functions of the ESCRT machinery in cell biology and viral replication. Biochem Soc Trans 45:613–634.

9. Flys T, Marlowe N, Hackett J, Parkin N, Schumaker M, Holzmayer V, Hay P, Eshleman SH. 2005. Analysis of PTAP duplications in the gag p6 region of subtype C HIV type 1. AIDS Res Hum Retroviruses 21:739–741.

10. Sharma S, Aralaguppe SG, Abrahams MR, Williamson C, Gray C, Balakrishnan P, Saravanan S, Murugavel KG, Solomon S, Ranga U. 2017. The PTAP sequence duplication in HIV-1 subtype C Gag p6 in drug-naive subjects of India and South Africa. BMC Infect Dis 17:95.

11. Garrus JE, von Schwedler UK, Pornillos OW, Morham SG, Zavitz KH, Wang HE, Wettstein DA, Stray KM, Cote M, Rich RL, Myszka DG, Sundquist WI. 2001. Tsg101 and the vacuolar protein sorting pathway are essential for HIV-1 budding. Cell 107:55–65.

12. Fisher RD, Chung HY, Zhai Q, Robinson H, Sundquist WI, Hill CP. 2007. Structural and biochemical studies of ALIX/AIP1 and its role in retrovirus budding. Cell 128:841–852.

13. Patil A, Bhattacharya J. 2012. Natural deletion of L35Y36 in p6 gag eliminate LYPXnL/ALIX auxiliary virus release pathway in HIV-1 subtype C. Virus Res 170:154–158.

14. Zhai Q, Landesman MB, Robinson H, Sundquist WI, Hill CP. 2011. Identification and structural characterization of the ALIX-binding late domains of simian immunodeficiency virus SIVmac239 and SIVagmTan-1. J Virol 85:632–637.

15. Neogi U, Rao SD, Bontell I, Verheyen J, Rao VR, Gore SC, Soni N, Shet A, Schulter E, Ekstrand ML, Wondwossen A, Kaiser R, Madhusudhan MS, Prasad VR, Sonnerborg A. 2014. Novel tetra-peptide insertion in Gag-p6 ALIX-binding motif in HIV-1 subtype C associated with protease inhibitor failure in Indian patients. Aids 28:2319–2322.

16. Neogi U, Engelbrecht S, Claassen M, Jacobs GB, van Zyl G, Preiser W, Sonnerborg A. 2016. Mutational Heterogeneity in p6 Gag Late Assembly (L) Domains in HIV-1 Subtype C Viruses from South Africa. AIDS Res Hum Retroviruses 32:80–84.

17. Aralaguppe SG, Winner D, Singh K, Sarafianos SG, Quinones-Mateu ME, Sonnerborg A, Neogi U. 2017. Increased replication capacity following evolution of PYxE insertion in Gag-p6 is associated with enhanced virulence in HIV-1 subtype C from East Africa. J Med Virol 89:106–111.

18. Kiguoya MW, Mann JK, Chopera D, Gounder K, Lee GQ, Hunt PW, Martin JN, Ball TB, Kimani J, Brumme ZL, Brockman MA, Ndung’u T. 2017. Subtype-Specific Differences in Gag-Protease-Driven Replication Capacity Are Consistent with Intersubtype Differences in HIV-1 Disease Progression. J Virol 91.

19. Chaturbhuj D, Patil A, Gangakhedkar R. 2018. PYRE insertion within HIV-1 subtype C p6-Gag functions as an ALIX-dependent late domain. Sci Rep 8:8917.

20. Neogi U, Singh K, Aralaguppe SG, Rogers LC, Njenda DT, Sarafianos SG, Hejdeman B, Sonnerborg A. 2018. Ex-vivo antiretroviral potency of newer integrase strand transfer inhibitors cabotegravir and bictegravir in HIV type 1 non-B subtypes. Aids 32:469–476.

21. Zhai Q, Fisher RD, Chung HY, Myszka DG, Sundquist WI, Hill CP. 2008. Structural and functional studies of ALIX interactions with YPX(n)L late domains of HIV-1 and EIAV. Nat Struct Mol Biol 15:43–49.

22. Sundquist WI, Schubert HL, Kelly BN, Hill GC, Holton JM, Hill CP. 2004. Ubiquitin recognition by the human TSG101 protein. Mol Cell 13:783–789.

23. Schindelin J, Arganda-Carreras I, Frise E, Kaynig V, Longair M, Pietzsch T, Preibisch S, Rueden C, Saalfeld S, Schmid B, Tinevez JY, White DJ, Hartenstein V, Eliceiri K, Tomancak P, Cardona A. 2012. Fiji: an open-source platform for biological-image analysis. Nat Methods 9:676–682.

24. Sette P, Nagashima K, Piper RC, Bouamr F. 2013. Ubiquitin conjugation to Gag is essential for ESCRT-mediated HIV-1 budding. Retrovirology 10:79.

25. Sharma S, Arunachalam PS, Menon M, Ragupathy V, Satya RV, Jebaraj J, Ganeshappa Aralaguppe S, Rao C, Pal S, Saravanan S, Murugavel KG, Balakrishnan P, Solomon S, Hewlett I, Ranga U. 2018. PTAP motif duplication in the p6 Gag protein confers a replication advantage on HIV-1 subtype C. J Biol Chem doi:10.1074/jbc.M117.815829.

26. Martins AN, Waheed AA, Ablan SD, Huang W, Newton A, Petropoulos CJ, Brindeiro RD, Freed EO. 2016. Elucidation of the Molecular Mechanism Driving Duplication of the HIV-1 PTAP Late Domain. J Virol 90:768–779.

27. Wright JK, Brumme ZL, Carlson JM, Heckerman D, Kadie CM, Brumme CJ, Wang B, Losina E, Miura T, Chonco F, van der Stok M, Mncube Z, Bishop K, Goulder PJ, Walker BD, Brockman MA, Ndung’u T. 2010. Gag-protease-mediated replication capacity in HIV-1 subtype C chronic infection: associations with HLA type and clinical parameters. J Virol 84:10820–10831.

28. Singh K, Flores JA, Kirby KA, Neogi U, Sonnerborg A, Hachiya A, Das K, Arnold E, McArthur C, Parniak M, Sarafianos SG. 2014. Drug resistance in non-B subtype HIV-1: impact of HIV-1 reverse transcriptase inhibitors. Viruses 6:3535–3562.

29. Bello NF, Wu F, Sette P, Dussupt V, Hirsch VM, Bouamr F. 2011. Distal leucines are key functional determinants of Alix-binding simian immunodeficiency virus SIV(smE543) and SIV(mac239) type 3 L domains. J Virol 85:11532–11537.

30. Amogne W, Bontell I, Grossmann S, Aderaye G, Lindquist L, Sonnerborg A, Neogi U. 2016. Phylogenetic Analysis of Ethiopian HIV-1 Subtype C Near Full-Length Genomes Reveals High Intrasubtype Diversity and a Strong Geographical Cluster. AIDS Res Hum Retroviruses 32:471–474.

31. Tully DC, Wood C. 2010. Chronology and evolution of the HIV-1 subtype C epidemic in Ethiopia. Aids 24:1577–1582.

32. Demirov DG, Orenstein JM, Freed EO. 2002. The late domain of human immunodeficiency virus type 1 p6 promotes virus release in a cell type-dependent manner. J Virol 76:105–117.

33. Huang M, Orenstein JM, Martin MA, Freed EO. 1995. p6Gag is required for particle production from full-length human immunodeficiency virus type 1 molecular clones expressing protease. J Virol 69:6810–6818.

34. Strack B, Calistri A, Craig S, Popova E, Gottlinger HG. 2003. AIP1/ALIX is a binding partner for HIV-1 p6 and EIAV p9 functioning in virus budding. Cell 114:689–699.

35. Dussupt V, Javid MP, Abou-Jaoude G, Jadwin JA, de La Cruz J, Nagashima K, Bouamr F. 2009. The nucleocapsid region of HIV-1 Gag cooperates with the PTAP and LYPXnL late domains to recruit the cellular machinery necessary for viral budding. PLoS Pathog 5:e1000339.

36. Gottwein E, Jager S, Habermann A, Krausslich HG. 2006. Cumulative mutations of ubiquitin acceptor sites in human immunodeficiency virus type 1 gag cause a late budding defect. J Virol 80:6267–6275.

37. Zhadina M, McClure MO, Johnson MC, Bieniasz PD. 2007. Ubiquitin-dependent virus particle budding without viral protein ubiquitination. Proc Natl Acad Sci U S A 104:20031–20036.

38. Coutsinos D, Invernizzi CF, Moisi D, Oliveira M, Martinez-Cajas JL, Brenner BG, Wainberg MA. 2011. A template-dependent dislocation mechanism potentiates K65R reverse transcriptase mutation development in subtype C variants of HIV-1. PLoS One 6:e20208.

39. Coutsinos D, Invernizzi CF, Xu H, Moisi D, Oliveira M, Brenner BG, Wainberg MA. 2009. Template usage is responsible for the preferential acquisition of the K65R reverse transcriptase mutation in subtype C variants of human immunodeficiency virus type 1. J Virol 83:2029–2033.

